# Multilevel Characterization Reveals Divergent Heat Responses Among Diploid, Tetraploid, and Hexaploid Wheat Genotypes

**DOI:** 10.64898/2026.01.22.701169

**Authors:** Anita Arenas-M, Ignacio Miño, Diego Landaeta, Cristóbal Uauy, Daniel F. Calderini, Javier Canales

## Abstract

Understanding heat stress (HS) responses in wheat is important for breeding varieties adapted to increasing temperature. HS during grain filling reduces wheat yield, but transcriptional and alternative-splicing responses among wheat genotypes of different ploidy remain poorly characterized. We conducted two field experiments with diploid *Triticum monococcum*, tetraploid *T. turgidum* subsp. *durum*, and hexaploid *T. aestivum*. In Exp. 2, one genotype per species was exposed to severe HS for four consecutive days beginning 10 days after anthesis, and grains were sampled for RNA sequencing 3 h after treatment onset. Mean grain yield declined by 59.4%, 19.0%, and 37.9% in the diploid, tetraploid, and hexaploid genotypes, respectively, accompanied by reductions in thousand-grain weight. After homoeolog grouping, the tetraploid genotype had the smallest fraction of heat-responsive groups and the hexaploid the largest. HS also altered alternative splicing, with broader functional enrichment in the hexaploid genotype. In this genotype, all three homoeologs of an *NF-YB* gene showed heat-induced skipping of exon E6, and the splice products of the D-genome locus were confirmed by RT-PCR and Sanger sequencing. These results show that the three examined genotypes differed in phenotypic, transcriptional, and splicing responses to HS and identify *NF-YB* splicing as a candidate component of the hexaploid grain response.

**Main conclusion:** Early grain-filling heating reduced yield and altered transcription and splicing in the three examined wheat genotypes, including increased *NF-YB* exon E6 skipping in the hexaploid wheat genotype.

## 1 Introduction

Wheat (*Triticum aestivum* L.) is a major global staple that supplies roughly 20% of human dietary calories and occupies about one-sixth of cultivated land (Guarin et al. 2022). Its broad agro-ecological adaptability, together with the unique viscoelastic properties of wheat flour, has enabled worldwide cultivation (Shiferaw et al. 2013). Despite advances in breeding and agronomy, current productivity gains may not meet projected demand increases of 35–56% by 2050 relative to 2010 (van Dijk et al. 2021). The success of wheat as a global crop is intrinsically linked to its polyploid nature, which arose through hybridization between different species and subsequent genome duplications. The presence of multiple subgenomes (A, B, and D) in modern hexaploid bread wheat (*T. aestivum* L., AABBDD genome) has enhanced its adaptive plasticity and contributed to its domestication and cultivation across diverse environments (Lv et al. 2021). Cultivated wheat species exhibit a range of ploidy levels, from ancestral diploids to modern tetraploid durum wheat (*T. turgidum* subsp. *durum*) and hexaploid bread wheat (*T. aestivum* L.). These successive hybridization events are evolutionarily recent: tetraploid wheat arose less than 500,000 years ago, and hexaploid wheat emerged less than 10,000 years ago (Krasileva et al. 2017). This genetic redundancy has allowed wheat to tolerate high densities of mutations, which can be exploited to uncover hidden variation, particularly in stress responses (Krasileva et al. 2017). Furthermore, increased genome and cell size associated with polyploidy has been hypothesized to buffer metabolic fluctuations and stabilize cellular processes under stress (Tossi et al. 2022). Although polyploidy may confer distinct adaptive capabilities and stress responses, systematic comparative studies across *Triticum* species with different ploidy levels remain scarce.

Heat stress (HS), particularly during critical developmental stages, severely impacts wheat productivity by disrupting key physiological processes and reducing grain yield potential (Akter and Rafiqul Islam 2017; Farooq et al. 2011). Recent studies indicate that wheat yields decline by approximately 6% for each degree Celsius increase in temperature (Zhao et al. 2017), and terminal HS can cause yield reductions of 15–16% in vulnerable regions such as South Asia and Africa (Pequeno et al. 2021). The detrimental effects of HS are most pronounced during reproductive and grain-filling phases (Farooq et al. 2011; Garćıa et al. 2015), leading to significant reductions in grain number, weight, and quality through impaired photosynthesis, accelerated senescence, and disrupted grain development (Farhad et al. 2023). Under high temperatures, wheat experiences metabolic limitations and oxidative damage to chloroplasts, which decreases dry matter accumulation and compromises grain yield (Farooq et al. 2011). Additionally, exposure to temperatures above 30*^◦^*C during floret formation can cause severe impairment of fertility. Temperatures above 34*^◦^*C during anthesis and grain filling reduce kernel weight, mainly by accelerating crop development, which in turn shortens the grain-filling period (Tack et al. 2015; Ńoia Júnior et al. 2023). As global temperatures continue to rise, understanding HS responses and developing adaptation strategies will be important for future wheat production and global food security.

Heat stress reduces kernel weight and impairs flour quality in wheat (Wardlaw et al. 2002) while altering regulation at transcriptional, post-transcriptional, and translational levels. Heat shock transcription factors (Hsfs) coordinate the transcriptional component of this response, and studies of individual Hsfs have identified regulatory mechanisms operating at both the protein and transcript levels. Temperature-sensitive SUMOylation of TaHsfA1 supports the early activation of downstream genes, whereas loss of this modification during sustained heat attenuates TaHsfA1 activity (Wang et al. 2023). At the transcript level, heat induces widespread changes in alternative splicing (AS) across the wheat transcriptome (Liu et al. 2018). For *TaHSFA6e*, the resulting heat-responsive splice isoforms differ in their ability to activate *TaHSP70* genes; TaHSP70 proteins subsequently promote stress-granule disassembly and translation reinitiation during recovery (Wen et al. 2023).

RNA sequencing analyses have revealed extensive transcriptome reprogramming in wheat grains exposed to high temperatures. Studies have shown that key heat response pathways in grains involve heat shock proteins (HSPs), chaperones, and genes related to protein folding, carbohydrate metabolism, and stress responses (Wang et al. 2019; Rangan et al. 2020; Arenas-M et al. 2021; Tomás et al. 2022; Paul et al. 2022). Transcriptomics of bread wheat exposed to heat found differential expression of genes associated with reactive oxygen species scavenging enzymes, hormone signaling, photosynthesis, and metabolic processes (Wang et al. 2019). Similarly, durum wheat grains exhibited upregulation of HSPs, antioxidant enzymes, and genes involved in protein processing, as well as downregulation of genes related to proteolysis and carbon metabolism under HS (Arenas-M et al. 2021). Traditional wheat landraces showed more extensive transcriptional changes compared to modern cultivars when subjected to heat, suggesting higher genetic diversity and altered regulation of stress response genes in landraces (Tomás et al. 2022). In hexaploid wheat, transcriptome dynamics also differed between grain and leaf tissues under HS (Rangan et al. 2020).

This study aimed to characterize HS responses in selected wheat genotypes representing three ploidy levels by combining phenotypic analyses with transcriptomic and alternative splicing studies. Specifically, we investigated: (1) the effects of HS on grain yield and quality traits in the diploid, tetraploid, and hexaploid genotypes, (2) the impact of HS on grain-filling dynamics and individual grain weight, (3) transcriptional responses through RNA-seq analysis, and (4) alternative splicing patterns.

## 2 Materials and methods

### 2.1 Plant Material and Field Experiments

Two field experiments were conducted at the Experimental Station of the Universidad Austral de Chile (UACh) in Valdivia (39*^◦^*47*^′^*S, 73*^◦^*14*^′^* W, 19 m asl), Chile, during the 2016/17 (Exp. 1) and 2017/18 (Exp. 2) growing seasons. The experiments were sown on August 11, 2016, and September 10, 2017, respectively. Thermal treatments were applied during the early grain-filling period (10–14 days after anthesis, DAA) for four consecutive days. The timing and duration of HS treatments were based on (i) the expected occurrence of heat waves in southern Chile and (ii) previous studies reporting high wheat sensitivity to temperature during early grain filling (Stone and Nicolas 1994; Lizana and Calderini 2013).

In Exp. 1, two genotypes per ploidy level were evaluated. The diploid *Triticum monococcum* lines CWI16964 (Mon1) and CWI19205 (Mon8) were obtained from CIM-MYT. The tetraploid *Triticum turgidum* subsp. *durum* cultivars were Corcolen-INIA (Corcolen) and Queule-INIA (Queule), and the hexaploid *Triticum aestivum* cultivars were Fritz-Baer (Fritz) and Impulso-Baer (Impulso). Diploid genotypes were sown 21 days earlier than tetraploids and hexaploids to synchronize anthesis (Zadoks 65) and physiological maturity (Zadoks 95). In Exp. 2, the genotype with the highest grain yield under control conditions in Exp. 1 was selected from each ploidy level: Mon8, Queule, and Fritz. Thus, Exp. 2 compared one yield-selected genotype from each ploidy level.

Thermal treatments in Exp. 1 comprised an ambient control (CT) and a moderate heating regime (TT1). Exp. 2 included CT and TT1 together with a severe heating regime (TT2). These labels describe treatment categories, not fixed temperature increments. Across entries, four-day mean temperatures were 22.7–28.5*^◦^*C for TT1 and 31.2–33.7*^◦^*C for TT2; the corresponding increases over the paired controls were 7.0–15.0*^◦^*C and 13.5–18.8*^◦^*C, respectively (Online Resource 5; Supplementary Table S1). Both experiments followed a randomized complete block design with three blocks. The plot was the experimental unit, and genotype × thermal-treatment combinations were randomized within each block. Plots were 1.4 m long with seven rows spaced 0.15 m apart, flanked by border rows of the spring wheat cultivar “Pantera-INIA.” Before sowing, 7 Mg ha*^−^*^1^ of CaCO_3_ was applied to reduce aluminum toxicity in the acidic soils. Plots were fertilized with 150 kg ha*^−^*^1^ of N and 300 kg ha*^−^*^1^ of P_2_O_5_ at sowing, with an additional 150 kg ha*^−^*^1^ of N applied during mid-tillering. Irrigation, pest control, and weed management were performed as needed.

### 2.2 Chambers and Devices for Temperature Control

Transparent polyethylene chambers (2.0 × 1.8 × 1.5 m) were constructed to increase temperature during grain filling. Each heated plot was enclosed by an independently controlled chamber, giving one chamber per heated treatment × genotype combination in each block (three chamber replicates per combination). Ambient-control plots were unchambered. Chambers remained on their assigned plots throughout the four-day exposure. Each chamber was equipped with thermostatically controlled electric heaters and thermal sensors placed at spike height and connected to a temperature regulator (Cavadevices, Buenos Aires, Argentina). Air temperature at spike height was recorded inside and outside the chambers using data loggers (Extech Instruments SDL200, FLIR Systems, Inc.). The values in Online Resource 5 (Supplementary Table S1) are means over the complete four-day exposure. Ambient temperature, precipitation, and photosynthetically active radiation (PAR) were monitored by the “Austral” weather station (INIA), located 50 m from the experimental site.

### 2.3 Plant Measurements

Crop phenology was monitored every two days starting at heading using the Zadoks scale (Zadoks et al. 1974). At anthesis (DC 65), 40 spikes per plot were tagged for subsequent measurements. Above-ground biomass was harvested at maturity from a 1-m section of the central row. Biomass samples were divided into spikes, leaf blades, and stems plus sheath leaves, dried at 65*^◦^*C for 48 hours, and weighed. Grain yield (GY), harvest index (HI), thousand-grain weight (TGW), and grain number (GN) were determined. Individual grain weight (IGW) was measured for grains at position 1 (G1) from two central spikelets of 10 spikes per plot. Grain dimensions (length, width, and area) were recorded using a Marvin seed analyzer (Wittenburg, Germany), and dry weights were obtained after drying at 65*^◦^*C for 48 hours.

For analysis of grain-filling dynamics, two spikes per plot were sampled twice weekly from anthesis until 52 or 60 days after anthesis, depending on genotype. Grains at position G1 were collected from the two central spikelets, dried at 65 *^◦^*C for 48 h, and weighed. Grain-filling rate and duration were estimated from the resulting dry-mass time courses using a bilinear model. Grain-filling duration and rate were calculated using a bilinear model (Calderini et al. 1999), with physiological maturity (PM) estimated using Table Curve V 3.0 (Jandel 1991). Thermal exposure was also expressed as heat load, calculated over the four treatment days as cumulative degree days above a base temperature of 18*^◦^*C.

### 2.4 Grain Protein and Starch Content

Total nitrogen (N) concentration in grains was determined using the Kjeldahl method. Protein concentration was calculated as N × 5.7 and expressed as a percentage of dry weight (Merrill and Watt 1973). Starch concentration, also expressed as a percentage of dry weight, was quantified using a Starch Colorimetric/Fluorometric Assay Kit (BioVision, Inc., USA). Briefly, 10 mg of wheat flour was washed with 90% ethanol, and starch was extracted using 10 N KOH and neutralized with 10 M H_3_PO_4_. Absorbance was measured at 570 nm using a Nanoquant Infinite M200 spectrophotometer (Tecan, Mannedorf, Switzerland).

### 2.5 RNA Isolation and Quantitative Real-Time PCR Analysis

For each biological replicate, complete caryopses were harvested from the two basal grains of four central spikelets on the main spike of each of five plants and pooled. Samples were immediately flash-frozen in liquid nitrogen and stored at −80*^◦^*C until extraction. Three independent biological replicates were collected for each experimental condition.

Total RNA was isolated following a modified CTAB-based protocol as described in Arenas-M et al. (2021) (Arenas-M et al. 2021). RNA quality and concentration were assessed using a Nanoquant Infinite M200 spectrophotometer (Tecan, Mannedorf, Switzerland), with samples exhibiting A260/A280 ratios between 1.8–2.0 considered suitable for downstream applications.

For cDNA synthesis, 500 ng of high-quality total RNA was reverse-transcribed using the 5X All-In-One RT MasterMix (Applied Biological Materials Inc., Vancouver, Canada) according to the manufacturer’s instructions. Gene expression was quantified using touchdown quantitative PCR (qPCR) assays as described by Zhang et al. (2015) (Zhang et al. 2015) to improve specificity and sensitivity. Reactions were performed using Brilliant II SYBR Green QPCR Master Mix (Agilent Technologies, Inc., Santa Clara, CA, USA) on an AriaMx Real-Time PCR System (Agilent Technologies).

Each qPCR reaction contained 25 ng of cDNA template and 200 nM of each gene-specific primer in a final volume of 25 *µ*L. The thermal cycling profile consisted of initial denaturation at 95*^◦^*C for 10 min; three touchdown cycles (95*^◦^*C for 20 s and 66*^◦^*C for 10 s, decreasing by 3*^◦^*C per cycle); and 40 amplification cycles (95*^◦^*C for 20 s, 55*^◦^*C for 10 s, and 72*^◦^*C for 10 s). Gene-specific forward and reverse primers are detailed in Online Resource 11 (Supplementary Table S7).

Raw fluorescence data were analyzed using Real-time PCR Miner version 4.0, which estimates threshold cycle (*C_t_*) values and reaction-specific amplification efficiencies from individual amplification curves (Zhao and Fernald 2005). The gene encoding a putative ubiquitin-conjugating enzyme (*TraesCS4A02G414200* ), previously validated for stable expression during wheat seed development (Borrill et al. 2016), was used as the internal reference. Relative expression was obtained by normalizing the efficiency-corrected target-gene estimates to the reference-gene estimates and expressing these ratios relative to the control, following the procedure of Zhao and Fernald (2005).

### 2.6 RNA-Seq Data Analysis

RNA sequencing was performed on grain samples collected 3 h after imposing HS during early grain filling (10 days after anthesis) under field conditions. Three biological replicates were sequenced for each genotype × condition combination (18 libraries). Samples were immediately flash-frozen in liquid nitrogen after field collection and stored at −80*^◦^*C until shipment. The samples were then transported to Novogene (Sacramento, CA, USA), where RNA quality assessment, library preparation, and sequencing were performed. For each sample, an Illumina RNA-seq library was constructed using the Illumina TruSeq Stranded Total RNA Sample Prep Kit according to the manufacturer’s protocol. Unique barcodes were incorporated during library preparation. Sequencing was performed on an Illumina NovaSeq 6000 platform using 2 × 150-bp paired-end reads, with a target depth of approximately 7–8 Gb per sample. Average Q20 and Q30 scores exceeded 97.4% and 93.8%, respectively. The mean clean-read retention rate (clean reads/raw reads) was 97.74%, and mean GC content ranged from 53.91% to 58.03% (Online Resource 12; Supplementary Table S8).

Raw sequencing reads were pseudo-aligned and quantified using Kallisto (v0.50.1) (Bray et al. 2016), a lightweight tool for accurate transcript-level abundance estimation via k-mer matching. A Kallisto index was first constructed from the reference transcriptomes of *Triticum aestivum* (IWGSC RefSeq v2.1) (Zhu et al. 2021), *Triticum turgidum* (Svevo v1) (Maccaferri et al. 2019), and *Triticum monococcum* (TA10622 v1.1) (Ahmed et al. 2023). This step employed the kallisto index command with a default k-mer size of 31. Transcript abundances were then quantified in paired-end mode using kallisto quant with default parameters.

Differential expression analysis was conducted separately within each genotype using DESeq2 v1.42 (Love et al. 2014) in R, with three biological replicates per condition. Transcript-level quantifications were summarized to gene-level counts using tximport (Soneson et al. 2015). A gene was considered differentially expressed at adjusted *p*-value *<* 0.05 (Benjamini–Hochberg correction) and absolute log_2_ fold change *>* 0.58. The design matrix included condition (HS versus control) as the factor of interest. The respective analyses tested 18,223, 39,847, and 68,303 expressed genes in Mon8, Queule, and Fritz. Homoeolog grouping reduced these sets to 18,223, 22,092, and 26,268 nonredundant gene units, respectively.

PCA was performed on normalized transcripts-per-million (TPM) values using scikit-learn (Pedregosa et al. 2011) in Python to assess sample clustering and variability. Data were standardized using z-score normalization prior to PCA computation. Confidence ellipses in the PCA plot were calculated using a 90% confidence interval.

Visualization of differential expression results was performed using volcano plots, generated with Matplotlib v3.7.1 in Python. The volcano plots were annotated with gene names for significantly differentially expressed genes, with upregulated genes shown in red and downregulated genes in blue.

GO enrichment analysis was performed using the clusterProfiler package (Xu et al. 2024) with GO annotations from the Triticeae-GeneTribe database (Chen et al. 2020). Upregulated and downregulated genes were analyzed separately using a Benjamini–Hochberg-adjusted *p*-value threshold of 0.05. Redundant terms were reduced by semantic similarity using rrvgo (Sayols 2023).

### 2.7 Alternative Splicing Analysis

Alternative splicing events were identified and analyzed using the Whippet software v1.6 (Sterne-Weiler et al. 2018), which enables efficient quantification of splicing events from RNA-seq data. Whippet indices were generated for each genome using corresponding GTF annotations via whippet-index.jl script (Sterne-Weiler et al. 2018). Prior to index generation, GFF3 files were converted to GTF format using AGAT (v1.3.2) (Dainat et al. 2024).

RNA-seq data from control and heat-stressed samples, each with three biological replicates, were processed using Whippet v1.6 to quantify splicing events (Sterne-Weiler et al. 2018). The resulting output files contained percent spliced-in (PSI) values for each detected splicing event. For downstream statistical analysis and visualization, the betAS R package (Ascensão-Ferreira et al. 2024) was employed. The PSI data were imported using the getDataset function with the tool parameter set to “whippet”. Events were extracted with the getEvents function and filtered to include only high-confidence alternative splicing events using the filterEvents function. Only events classified as alternative acceptor (AA), alternative donor (AD), alternative first exon (AF), alternative last exon (AL), core exon (CE), retained intron (RI), tandem alternative polyadenylation site (TE), or tandem transcription start site (TS) were retained. Further filtering was performed with the alternativeEvents function to exclude constitutive events by retaining only those with PSI values between 5% and 95%.

Differential splicing analysis was conducted by comparing PSI values between control and heat-stressed samples using betAS (Ascensão-Ferreira et al. 2024). Statistical significance was assessed using the beta distribution-based approach implemented in prepareTableVolcanoFDR, with 1,000 simulations and FDR correction. Events were considered differential at FDR *<* 0.05 and |ΔPSI| *>* 0.1. For GO analysis, events were separated by the sign of ΔPSI before collapsing to unique gene identifiers; consequently, a gene containing both increased- and decreased-PSI events was included in both gene sets.

Individual splicing events of interest were visualized using the plotIndividualDen-sitiesList function from the betAS package (Ascensão-Ferreira et al. 2024), which displays the density distributions of PSI values for each condition. Additionally, sashimi plots were generated using ggsashimi (Garrido-Martın et al. 2018) to visualize read coverage and splicing patterns across exon-intron junctions for selected events.

Selected alternative splicing events were validated using RT-PCR. Total RNA was extracted from wheat grains using the Plant Total RNA Mini Kit (Geneaid, RPD100). First-strand cDNA was synthesized from 1.0 *µ*g RNA using All-In-One 5X RT Mas-terMix (abm, G592). PCR amplification was performed with GoTaq G2 Flexi DNA Polymerase for 30 cycles. Products were separated on 2.5% agarose gels at 70 V for 90 min, and band intensities were quantified using GelAnalyzer 23.1.1. E6 inclusion was calculated as the intensity of the E6-included band divided by the sum of the E6-included and E6-skipped bands. Control and HS values from three paired biological replicates were compared by a two-sided paired *t*-test. For sequence confirmation, PCR products were cloned into pJET1.2 using the CloneJET PCR Cloning Kit, transformed into One Shot TOP10 competent *E. coli*, and sequenced by Sanger sequencing.

### 2.8 Statistical Analysis

The plot was the experimental unit for yield, yield-component, and grain-composition responses. Data were analyzed by two-way ANOVA with genotype and thermal treatment as fixed factors, followed by Tukey’s multiple-comparisons test (GraphPad Prism 8.1.1); block was not included in the reported model. The displayed SEM is the pooled residual standard error from this model, with three plot replicates per genotype × treatment combination. Linear regression analyses were used to assess relationships between variables. RNA-seq, alternative-splicing, RT-qPCR, and RT-PCR analyses used the biological replication and inferential procedures specified in their respective subsections. Significance levels for the field-trait analyses are indicated as \**p <* 0.05, \*\**p <* 0.01, \*\*\**p <* 0.001, and \*\*\*\**p <* 0.0001.

## 3 Results

### 3.1 Differential Effects of HS on Yield and Grain Quality in Diploid, Tetraploid, and Hexaploid Wheat Genotypes

Two independent field experiments were conducted in southern Chile to evaluate HS responses in wheat genotypes representing different ploidy levels. During the crop cycle, the average temperature was 13.0*^◦^*C in Experiment 1 (Exp. 1) and 13.4*^◦^*C in Experiment 2 (Exp. 2), and total rainfall was 618.7 mm and 498.6 mm, respectively (Online Resource 1; Supplementary Fig. S1). Rainfall, supplemental irrigation, and high soil water retention were intended to minimize water limitation. HS treatments were imposed for four consecutive days during early grain filling (10–14 days after anthesis) using temperature-controlled polyethylene chambers (Fig. 1). The treatments comprised an ambient control (CT) and a moderate heating regime (TT1) in both experiments; Exp. 2 also included a severe heating regime (TT2). TT1 and TT2 denote treatment categories rather than fixed temperature increments: mean temperatures measured at spike height over the four-day treatments ranged from 22.7 to 28.5*^◦^*C for TT1 and from 31.2 to 33.7*^◦^*C for TT2 (Online Resource 5; Supplementary Table S1). Exp. 1 revealed no significant difference (*p >* 0.05) in grain yield (GY) between the heat-stressed and control groups. In contrast, Exp. 2 showed a significant negative effect (*p <* 0.01) of HS on GY. GY reductions ranged from 9.7 to 59.4% relative to the control, with larger reductions under TT2 (Table 1). The genotype × treatment interaction was not significant (*p >* 0.05). In the diploid genotype, mean GY decreased by 36% under TT1 and 59.4% under TT2. The corresponding decreases were 9.7% and 19% in the tetraploid genotype and 12.3% and 37.8% in the hexaploid genotype (Table 1). Biomass, harvest index (HI), grain number (GN), and thousand-grain weight (TGW) showed similar treatment responses (Online Resource 6; Supplementary Table S2). TGW remained relatively stable under heating in Exp. 1; in Exp. 2, it decreased by 9.3–12.1% under TT1 and 14.4–23.2% under TT2 (Table 1).

**Fig. 1.**
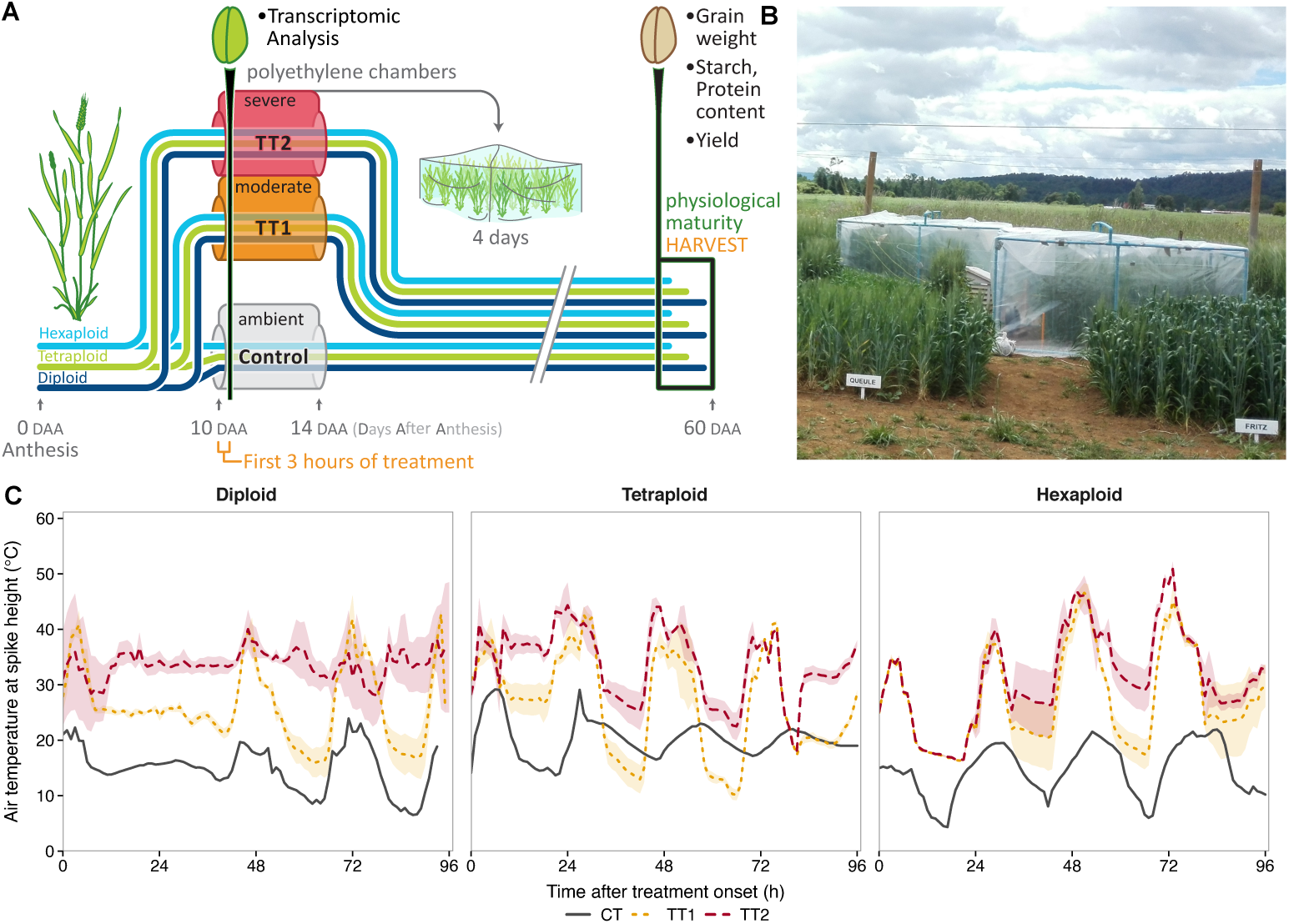
Experimental design and temperature profiles during the thermal treatments. (A) Schematic representation of the experimental setup. Diploid (*Triticum monococcum*), tetraploid (*T. turgidum* subsp. *durum*), and hexaploid (*T. aestivum*) wheat genotypes were exposed to an ambient control (CT), moderate heating (TT1), or severe heating (TT2; Exp. 2 only) beginning at 10 days after anthesis (DAA). Heated treatments were imposed using temperature-controlled polyethylene chambers, whereas CT plots remained unchambered. Developing grains were collected 3 h after treatment onset for transcriptomic analysis, and the thermal treatments continued for four consecutive days. Plants were harvested at physiological maturity for determination of yield, grain weight, and starch and protein concentrations. (B) Field implementation of the polyethylene chambers. (C) Air temperature measured at spike height during the four-day exposure in Exp. 2 for the diploid genotype Mon8, tetraploid genotype Queule, and hexaploid genotype Fritz. Lines represent hourly means, and shaded bands indicate *±* SEM across three replicate plots. TT1 and TT2 denote treatment categories rather than fixed temperature increments. Four-day mean temperatures and heat loads are reported in Online Resource 5 (Supplementary Table S1).

**Table 1.**
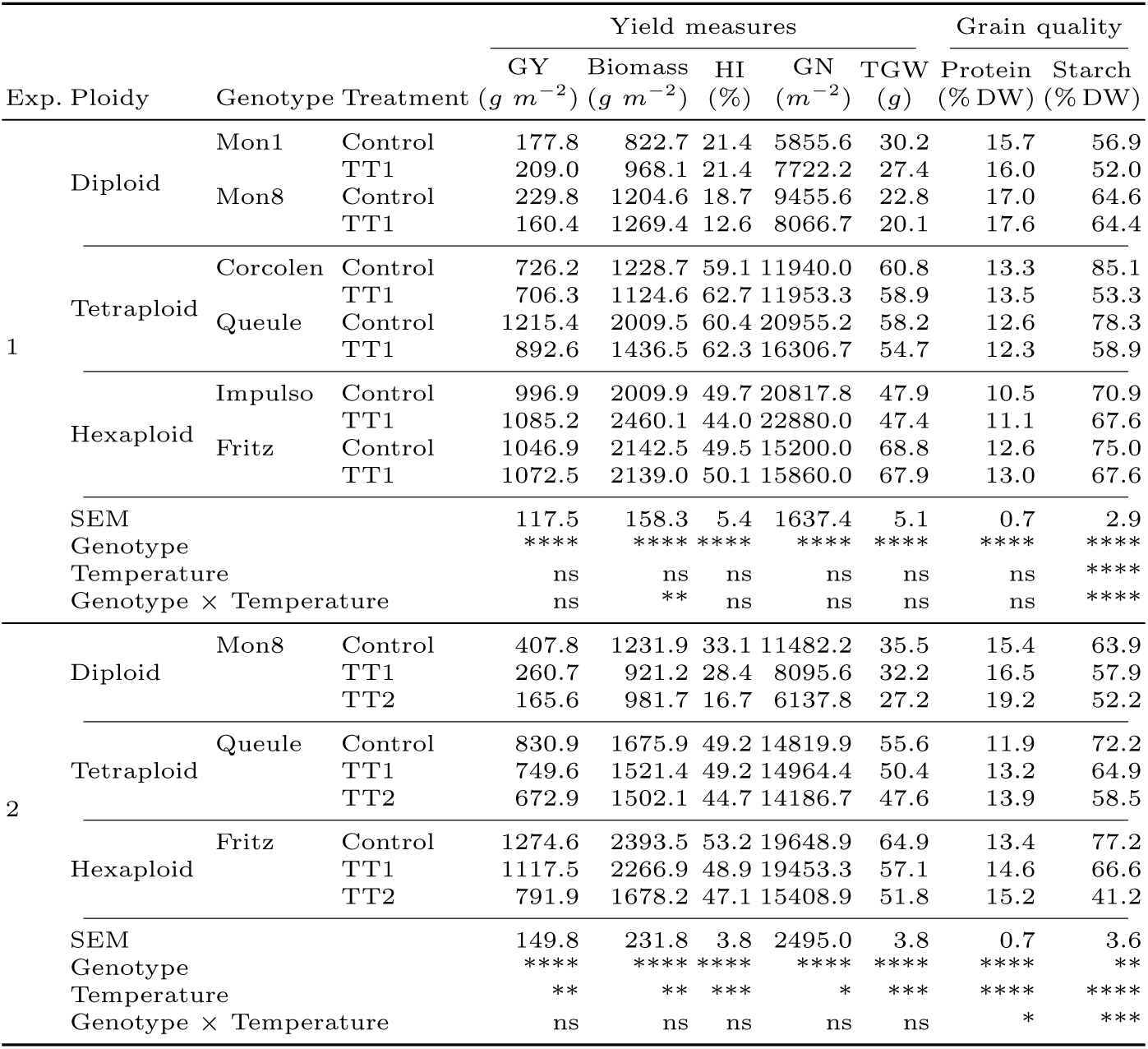
Effects of thermal treatments on yield components and grain quality in the examined wheat genotypes. The table presents data from two experiments comparing an ambient control (CT), moderate heating (TT1), and severe heating (TT2; Exp. 2 only) in diploid (*T. monococcum*), tetraploid (*T. turgidum* subsp. *durum*), and hexaploid (*T. aestivum*) wheat. GY, grain yield; HI, harvest index; GN, grain number; TGW, thousand-grain weight. Measured temperatures for each genotype and treatment are reported in Online Resource 5 (Supplementary Table S1). Starch and protein concentrations are expressed as percentages of dry weight (DW). The single SEM for each response is the pooled residual standard error from the ANOVA (*n* = 3 plot replicates per genotype *×* treatment combination). Statistical significance is indicated as ns (not significant), * (*P <* 0.05), ** (*P <* 0.01), *** (*P <* 0.001), and **** (*P <* 0.0001).

To characterize the effects of the heat treatments on grain quality, starch and protein concentrations were quantified at harvest. Under control conditions, diploid genotypes had the lowest grain starch concentrations (56.9–64.6% dry weight [DW]), whereas tetraploid and hexaploid cultivars had higher values (72.2–85.1% DW and 70.9–77.2% DW, respectively) (Table 1). Both experiments revealed significant differences in grain starch concentration among genotypes, with significant genotype × treatment interactions (Table 1). Across both experiments, TT1 produced average reductions in grain starch concentration of 6.0% in diploid and 9.3% in hexaploid genotypes. Tetraploid genotypes had a mean reduction of 24.0% under TT1 (Table 1). Under TT2 (Exp. 2), starch concentration decreased by 18.3%, 18.9%, and 46.6% in the diploid, tetraploid, and hexaploid genotypes, respectively (Table 1).

Under control conditions, diploid genotypes had grain protein concentrations of 15.4–17.0% DW, compared with 10.5–13.4% DW in the polyploid genotypes; the genotype effect was significant in both experiments (*p <* 0.0001; Table 1). TT1 had no significant effect on protein content in Exp. 1 (*p >* 0.05) and produced small increases in all three genotypes in Exp. 2 (Table 1).

### 3.2 Impact of HS on Grain-Filling Duration and Individual Grain Weight

Assessing the stability of grain weight formation across different ploidy levels provides information about wheat responses to rising temperatures (Li et al. 2018). Genotype and thermal treatments significantly influenced individual grain weight at position 1 (IGW-G1). In Experiment 1, moderate heat stress reduced IGW-G1 by 15.6–17.8% in the two diploid genotypes, whereas changes in the tetraploid and hexaploid genotypes ranged from a 3.5% increase to a 5.3% decrease. Under the severe thermal conditions of Experiment 2, IGW-G1 decreased by 46.7%, approximately 18%, and 21% in the diploid, tetraploid, and hexaploid genotypes, respectively (Online Resource 6; Supplementary Table S2).

IGW-G1 decreased with increasing mean temperature across the examined genotypes (Fig. 2A). In relative terms, the fitted reductions were 2.8% per *^◦^*C in the diploid genotype and 1.2–1.3% per *^◦^*C in the tetraploid and hexaploid genotypes (Fig. 2B). Grain-filling curves showed that HS shortened the filling period, particularly in the diploid genotype, where estimated duration decreased from 36.8 to 28.0 days, whereas the tetraploid genotype maintained a duration of approximately 46 days across treatments (Fig. 2C; Online Resource 6, Supplementary Table S2). The shorter filling period coincided with the larger reduction in final grain weight.

**Fig. 2.**
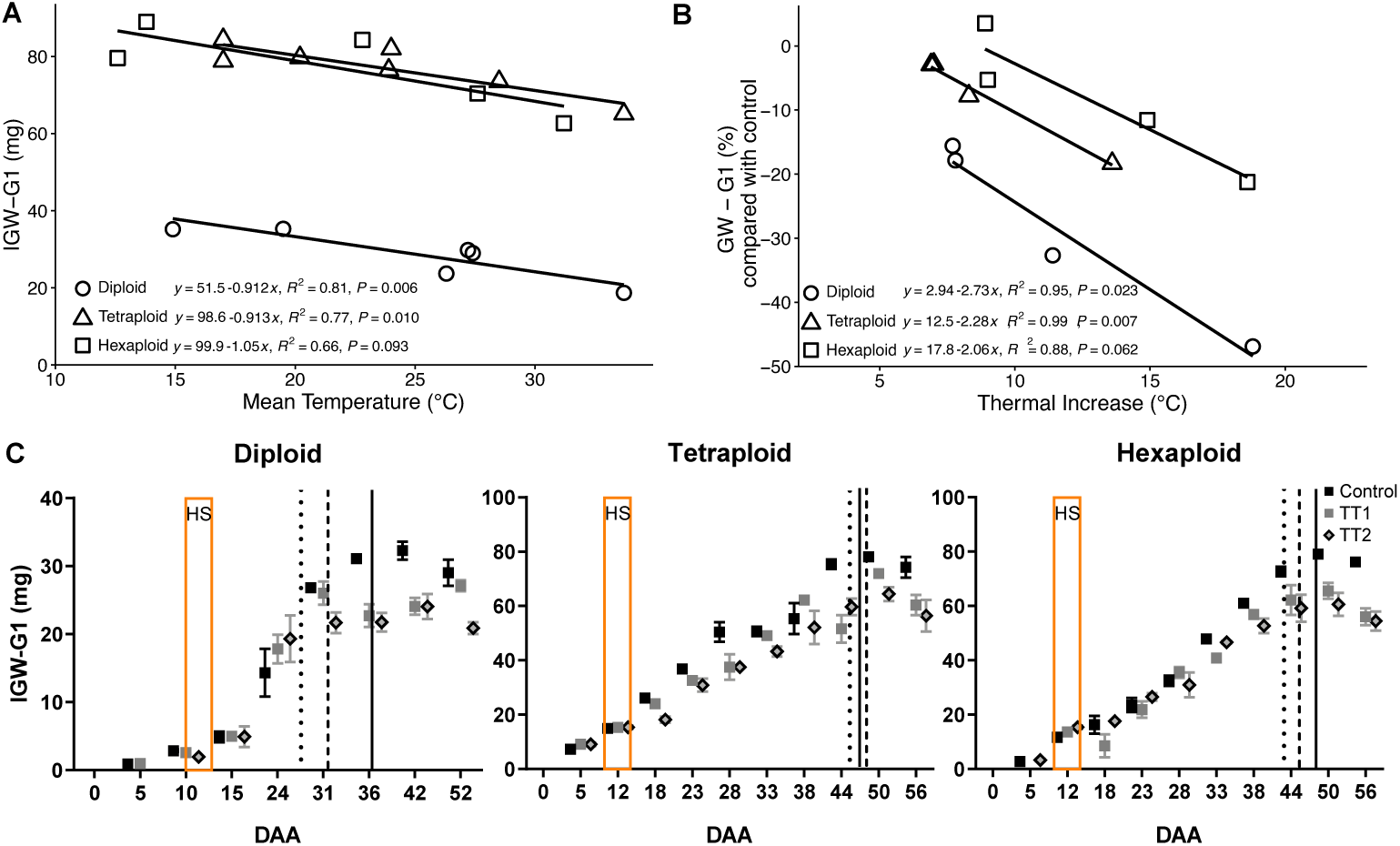
HS effects on grain weight during grain filling. (A) Relationship between mean temperature during the four-day treatment and individual grain weight at position 1 (IGW-G1) in the diploid, tetraploid, and hexaploid genotypes. Lines show fitted linear regressions; equations, coefficients of determination, and *P* -values are given in the panel. (B) Relative change in IGW-G1 from the corresponding ambient control plotted against the treatment-induced temperature increase. (C) Grain-filling time courses under CT, TT1, and TT2. Orange boxes indicate the four-day thermal treatments, and vertical lines mark the estimated treatment-specific timing of physiological maturity

### 3.3 Comparative RNA-seq Analysis of Heat Responses

Based on the yield and grain quality responses observed in our field experiments, we focused the RNA-seq analysis on the severe heat treatment (TT2) in Exp. 2, which produced the highest spike temperatures and the largest mean yield reductions (Online Resource 5; Supplementary Table S1). Mean yield declined by 59.4%, 19%, and 37.8% in the diploid, tetraploid, and hexaploid genotypes, respectively. Grain samples were collected 3 h after heating began at 10 DAA, when grain weight is particularly susceptible to HS damage (Farooq et al. 2011). Differential expression was tested separately for each genome using three biological replicates per condition.

Principal component analysis (PCA) separated control and heat-treated samples in all three analyses, indicating that treatment was a major source of transcriptional variation (Fig. 3A). The first principal component (PC1) accounted for the largest share of variance in each analysis, explaining 51.71%, 37.74%, and 43.21% in the diploid, tetraploid, and hexaploid genotypes, respectively (Fig. 3A).

**Fig. 3.**
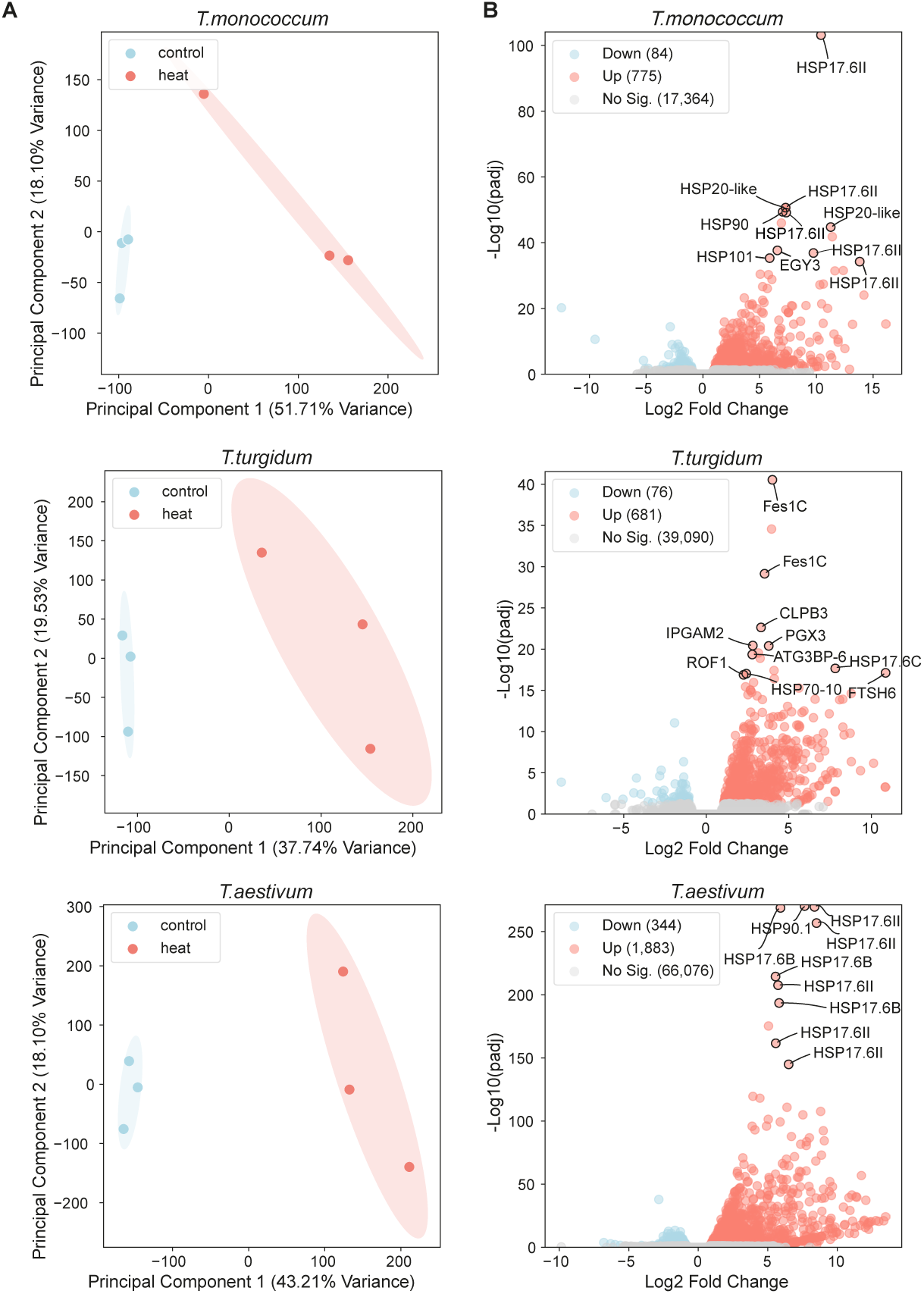
RNA-seq analysis of HS responses in the three examined wheat genotypes. (A) Principal component analysis of control (blue) and heat-treated (red) samples. Ellipses represent 90% confidence intervals. (B) Volcano plots of genome-specific differential expression analyses (adjusted *p*-value *<* 0.05 and *|* log_2_ fold change*| >* 0.58). Downregulated genes are blue, upregulated genes are red, and non-significant genes are grey; numbers in parentheses give the counts in each category. The y-axis is *−* log_10_ adjusted *p*-value and the x-axis is log_2_ fold change. The ten most significant DEGs are labeled. Plots were generated in Python using Matplotlib v3.7.1

The genome-specific analyses included 18,223, 39,847, and 68,303 expressed genes in the diploid, tetraploid, and hexaploid genotypes, respectively. After homoeolog grouping, these sets corresponded to 18,223, 22,092, and 26,268 nonredundant gene units, providing more comparable gene counts across ploidy levels. The diploid analysis identified 859 differentially expressed genes (DEGs), comprising 775 upregulated and 84 downregulated genes (| log_2_ FC| *>* 0.58, adjusted *p*-value *<* 0.05; Online Resource 7, Supplementary Table S3). Among these, several heat shock protein-encoding genes showed particularly strong induction (log_2_ FC *>* 10, − log_10_(*p_adj_*) *>* 40), including members of the HSP70 and HSP90 families.

The tetraploid analysis identified 757 DEGs (681 upregulated, 76 downregulated), corresponding to 562 homoeolog groups. Several molecular chaperone-encoding genes were strongly induced, including *Fes1C* (log_2_ FC = 4.0, − log_10_(*p_adj_*) *>* 40), *CLPB3* (log_2_ FC = 3.3, − log_10_(*p_adj_*) *>* 20), and *HSP17.6C* (log_2_ FC = 7.8, − log_10_(*p_adj_*) *>* 18).

The hexaploid analysis identified 2,227 DEGs (1,883 upregulated, 344 downregulated), representing 1,445 homoeolog groups. Gene expression changes were particularly pronounced for multiple *HSP17.6* family members and *HSP90.1*, which showed highly significant differential expression (− log_10_(*p_adj_*) *>* 200) and high fold changes (log_2_ FC *>* 7.5).

Comparative analysis of heat-responsive genes revealed both shared and contrasting patterns of molecular chaperone gene induction. *HSP17.6* genes were strongly induced in the diploid and hexaploid genotypes, whereas *Fes1C* and *CLPB3* were among the most strongly induced chaperone genes in the tetraploid genotype (Fig. 3B). To assess the concordance between RNA-seq and RT-qPCR, we quantified the expression of two selected heat-responsive genes, *HSP17.6II* (TraesCS3A03G024620) and *HSP26* (TraesCS4A03G0135000), in the three genotypes under control conditions and moderate (TT1) and severe (TT2) heat stress. RT-qPCR showed that both genes were significantly upregulated under heat stress relative to the control (*P <* 0.05), in agreement with the RNA-seq data (Online Resource 2; Supplementary Fig. S2). Expression of both genes increased from TT1 to TT2 in the tetraploid genotype. The diploid and hexaploid genotypes also showed strong induction under heat stress; however, *HSP26* induction was lower in the hexaploid than in the diploid and tetraploid genotypes. Overall, RT-qPCR reproduced the expression trends observed in the RNA-seq data for both genes.

### 3.4 Comparative GO Enrichment Analysis Identifies Shared and Genotype-Associated Heat Responses

To characterize the functions represented among the differentially expressed genes, we performed Gene Ontology (GO) enrichment analysis using clusterProfiler (Xu et al. 2024), followed by semantic similarity reduction with the rrvgo package (Sayols 2023) to obtain non-redundant GO terms (Online Resource 8; Supplementary Table S4). The analysis of upregulated genes (Fig. 4A) identified 14 non-redundant biological processes in the hexaploid, 12 in the diploid, and two in the tetraploid genotype. Heat response and protein folding terms (group I) were enriched in all three genotypes. The diploid and hexaploid genotypes shared nine GO terms (group II), including responses to abiotic stresses (salt, hypoxia, arsenic, and copper ion) and protein homeostasis (response to unfolded protein). Three terms were enriched in the hexaploid but not detected as significant in the other two analyses (group IV): response to molecules of fungal origin, protein quality control through the endoplasmic-reticulum-associated degradation (ERAD) pathway, and mRNA processing through cis splicing via the spliceosome.

**Fig. 4.**
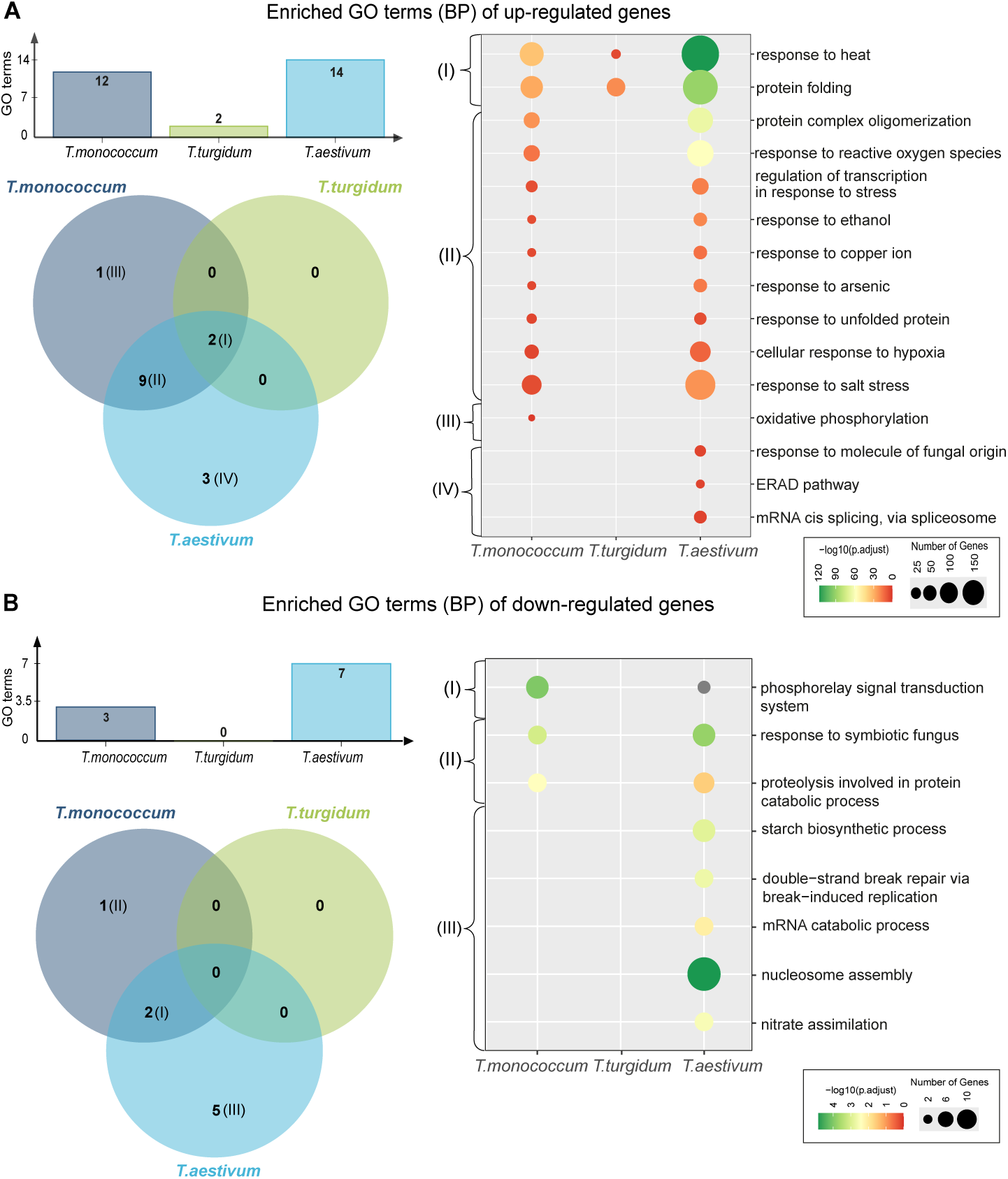
GO enrichment analysis of differentially expressed genes in the three examined wheat genotypes. (A) Enriched biological process terms among upregulated genes. The bar graph gives the number of enriched GO terms per genotype, the Venn diagram shows shared terms and terms detected in only one analysis, and the dot plot gives significance (*−* log_10_(*p*-adjust)) and gene count. (B) Corresponding analysis of downregulated genes. GO annotations were derived from *Triticeae-GeneTribe*(Chen et al. 2020); enrichment was tested using the clusterProfiler R package (Xu et al. 2024) with *p*-adjust *<* 0.05. P-values were adjusted by the Benjamini–Hochberg method. Roman numerals link Venn-diagram groups to the dot plots

The analysis of downregulated genes (Fig. 4B) identified seven non-redundant GO terms in the hexaploid, three in the diploid, and none in the tetraploid genotype. Five terms, including starch biosynthesis and mRNA catabolism, were enriched in the hexaploid but not detected as significant in the other two analyses (group III). Two GO terms were shared between the diploid and hexaploid genotypes (group I), involving phosphorelay signal transduction and responses to fungi.

These results indicate a shared core HS response, while the numbers and identities of enriched terms differed among the diploid, tetraploid, and hexaploid genotypes. The hexaploid genotype had the largest number of terms among both up- and downregulated genes.

### 3.5 Comparative Analysis of HS-Induced Alternative Splicing Across Wheat Species

GO enrichment analysis of upregulated genes in the hexaploid genotype identified the term “mRNA cis splicing, via spliceosome”, which was not significantly enriched in the other two genotypes (Fig. 4A). Based on this result and the known role of alternative splicing (AS) in stress responses, we hypothesized that AS patterns may differ among the diploid, tetraploid, and hexaploid genotypes. We therefore conducted a comparative analysis of AS events in response to HS across the three genotypes.

HS induced distinct patterns of AS among the examined genotypes (Fig. 5A; Online Resource 9, Supplementary Table S5). Differential events met both FDR *<* 0.05 and an absolute change in percent spliced in (|ΔPSI|) *>* 0.1. The diploid genotype had 116 events with increased PSI and 145 with decreased PSI; the tetraploid had 195 and 254, respectively; and the hexaploid had 159 and 208, respectively. Analysis of AS event types (Fig. 5B) indicated that alternative acceptor (AA) and core exon (CE) events were prevalent in all three genotypes, often followed by tandem alternative polyadenylation site (TE) or alternative donor (AD) events. AA events represented 35.4% of those in the tetraploid genotype, compared with 27.2% in the diploid and 27.8% in the hexaploid genotype. Retained intron (RI) events represented 12.0%, 11.1%, and 9.1% in the hexaploid, diploid, and tetraploid genotypes, respectively. GO enrichment of genes containing differential splicing events (Fig. 5C) yielded one term among diploid genes containing increased-PSI events (“embryo development ending in seed dormancy”). In the hexaploid genotype, genes containing decreased-PSI events were enriched for four terms related to mRNA splicing, nuclear protein import, chromatin organization, and regulation of gene expression, whereas genes containing increased-PSI events were enriched for two terms related to transcriptional regulation and peptidyl-serine dephosphorylation. No significant term was detected in the tetraploid genotype.

**Fig. 5.**
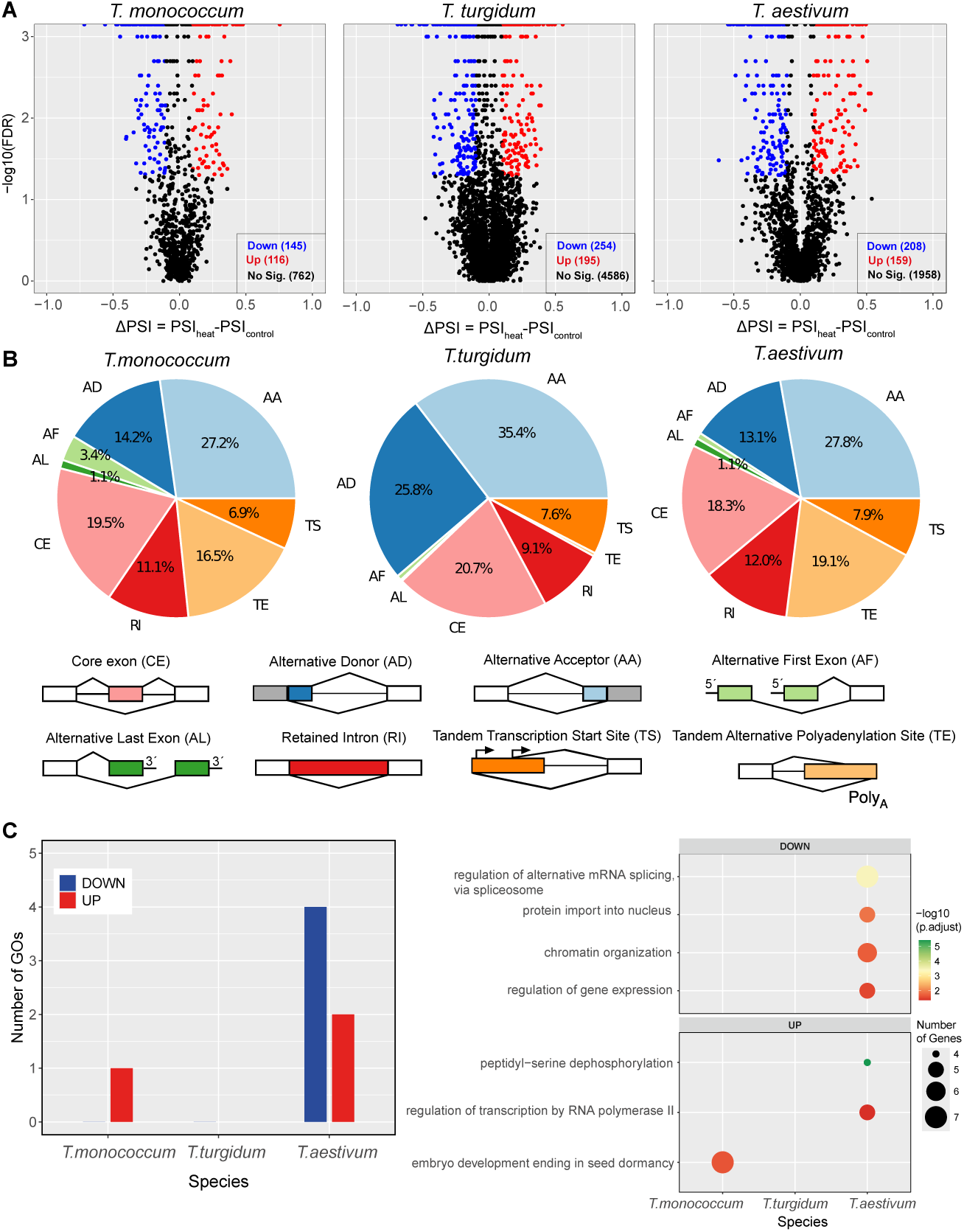
Alternative splicing events and GO enrichment under HS. (A) Volcano plots of ΔPSI = PSI_heat_ *−* PSI_control_. Blue and red dots represent events with significantly decreased and increased PSI, respectively (FDR *<* 0.05, *|*ΔPSI*| >* 0.1); black dots are non-significant. (B) Distribution of core exon (CE), alternative donor (AD), alternative acceptor (AA), alternative first exon (AF), alternative last exon (AL), retained intron (RI), tandem transcription start site (TS), and tandem alternative polyadenylation site (TE) events. (C) Number of enriched GO terms among genes containing decreased-PSI or increased-PSI events. Dot size indicates gene count and color gives *−* log_10_ adjusted *p*-value. Benjamini–Hochberg-adjusted *p*-values *<* 0.05 were considered significant

### 3.6 HS Induces Exon E6 Skipping in the Transcription Factor *NF-YB* in the Hexaploid Genotype

Given the enrichment of transcriptional regulation among alternatively spliced genes in *T. aestivum*, we focused on transcription factors whose splicing patterns might contribute to the HS response. Among the alternatively spliced transcription factors, *NF-YB* (*TraesCS1D03G0689500* ) emerged as a candidate for further analysis based on two criteria: (i) it showed significant heat-induced AS (|ΔPSI| *>* 0.1, FDR *<* 0.05) and (ii) the AS response was consistent across all three homoeologs (A, B, and D subgenomes; Online Resource 10, Supplementary Table S6). Exon E6 is defined relative to the *TraesCS1D03G0689500.1* transcript model. The concordant response of the three *NF-YB* homoeologs supported its selection for experimental analysis. Only the B-genome homoeolog of *T. turgidum* is shown in Fig. 6A because the A-genome homoeolog did not show a significant heat-induced change in PSI.

**Fig. 6.**
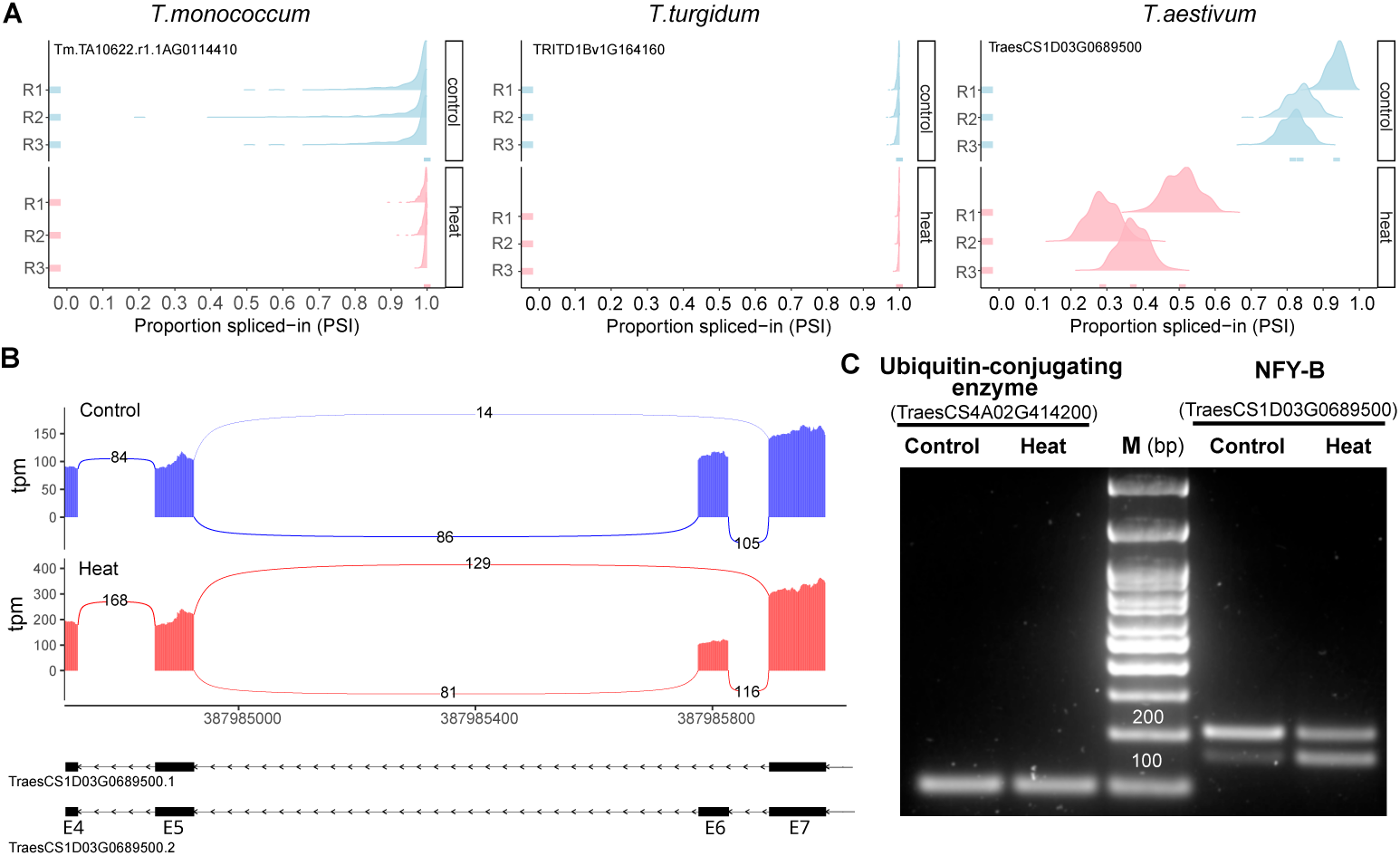
HS increases exon E6 skipping in *NF-YB* in the examined *Triticum aestivum* genotype. (A) PSI values across three biological replicates under control (blue) and HS (pink) conditions. Events are located at *T. aestivum* (*TraesCS1D03G0689500*, chr1D:387985776–387985826), *T. turgidum* (*TRITD1Bv1G164160*, chr1B:510670414–510670464), and *T. monococcum* (*Tm.TA10622.r1.1AG0114410*, chr1A:529294237–529294491). Only the *T. turgidum* B-genome homoeolog is shown because its A-genome homoeolog did not show a significant heat-induced change in PSI. Exon E6 refers to transcript model *TraesCS1D03G0689500.1*. (B) RNA-seq coverage and splice-junction counts for *TraesCS1D03G0689500* under control and HS. PSI was calculated from splice-junction reads using betAS (Ascensão-Ferreira et al. 2024); coverage is shown as transcripts per million (TPM). (C) RT-PCR detection of the approximately 200-bp E6-included amplicon and the approximately 150-bp E6-skipped amplicon. Ubiquitin-conjugating enzyme served as an amplification control. Amplicon identities were confirmed by cloning and Sanger sequencing

Analysis of *NF-YB* splicing patterns revealed significant, heat-induced alterations in the hexaploid genotype that were not detected in the diploid or tetraploid orthologs (Fig. 6). In the hexaploid genotype, the PSI for exon E6 decreased from 0.85 under control conditions to 0.37 after heat treatment (ΔPSI = −0.48), indicating a substantial increase in the proportion of transcripts skipping E6 (Fig. 6A).

Splice-junction counts changed in the same direction (Fig. 6B). Under control conditions the E5–E6 and E6–E7 junctions carried 86 and 105 reads while the E5–E7 junction that bypasses E6 carried 14; after heating the same junctions carried 81 and 116 reads against 129 for the skipping junction.

Reverse transcription polymerase chain reaction (RT-PCR) detected an approximately 200-bp E6-included amplicon (E5–E6–E7) and an approximately 150-bp E6-skipped amplicon (E5–E7) (Fig. 6C). Densitometry of three paired biological replicates gave E6 inclusion of 84.7 ± 3.4% in the control and 49.8 ± 2.7% under HS (mean ± SD; two-sided paired *t*-test, *t* = 28.6, *df* = 2, *p* = 0.0012; Online Resource 4, Supplementary Fig. S4). The control estimate matches the RNA-seq PSI (0.85), whereas the heated estimate is slightly higher than the sequencing value (0.50 versus 0.37).

To confirm the identity of the splice variants, the RT-PCR products were cloned and sequenced. Sanger sequencing verified that the ∼150-bp amplicon corresponds to the *NF-YB* isoform lacking exon E6, whereas the ∼200-bp amplicon corresponds to the E6-included isoform (Online Resource 3; Supplementary Fig. S3).

Collectively, these data indicate heat-induced alternative splicing of *NF-YB* in the hexaploid genotype, whereas the corresponding splicing pattern remained largely unchanged in the diploid and tetraploid genotypes under HS.

## 4 Discussion

Climate change presents significant challenges for global wheat production through increased weather variability and extreme conditions, particularly HS. This study evaluated HS responses during early grain filling in selected diploid, tetraploid, and hexaploid genotypes under field conditions. The three genotypes differed in mean yield loss and showed contrasting patterns of differential gene expression and AS. GO enrichment among genes containing differential splicing events yielded one term in the diploid and six in the hexaploid genotype. Heat-induced exon E6 skipping in *NF-YB* was experimentally supported in the hexaploid genotype.

### 4.1 HS Sensitivity of the Diploid, Tetraploid, and Hexaploid Genotypes

The contrasting yield responses between experiments indicate that the effects of short-term HS depended on treatment intensity and the prevailing thermal conditions (Wardlaw et al. 2002; Farooq et al. 2011). In Exp. 1, TT1 produced four-day mean temperatures of 22.7–27.4*^◦^*C, corresponding to increases of 7.0–9.0*^◦^*C above the paired ambient controls, and left grain yield and its components largely unaffected in the diploid, tetraploid, and hexaploid genotypes. Thus, a four-day heat episode did not invariably reduce yield under the conditions tested.

Exp. 2 showed yield reductions under both moderate (TT1) and severe (TT2) HS, with larger mean losses under TT2. Relative IGW-G1 decreased by 2.8% *^◦^*C*^−^*^1^ in the diploid genotype and by 1.2–1.3% *^◦^*C*^−^*^1^ in the tetraploid and hexaploid genotypes. Mean yield losses under TT2 were 59.4%, 19%, and 37.8%, respectively.

HS was associated with a shorter grain-filling duration. Under control conditions, diploid genotypes had shorter grain-filling periods (37–43 days) than tetraploid and hexaploid cultivars (46–49 days), and HS further shortened this developmental window. The relative stability of estimated grain-filling rates across thermal treatments is compatible with maintenance of assimilate supply during HS, potentially through continued remobilization of stem reserves (Tahir and Nakata 2005; Wang et al. 2018). Previous studies likewise found that heat can reduce grain weight mainly by truncating the grain-filling period while having little or no effect on mean filling rate (Zahedi and Jenner 2003; Shirdelmoghanloo et al. 2016). Together, these results suggest that shortening of the developmental window contributed to the final differences in grain weight.

Grain starch and protein concentrations also showed genotype-associated responses to HS. Under control conditions, diploid genotypes had higher grain protein concentrations (15.4–17.0% DW) than the polyploid varieties (10.5–13.4% DW). Starch accumulation was sensitive to HS. Under TT1, the tetraploid varieties had the largest mean decrease (24.0%), whereas under TT2 the hexaploid cultivar had a 46.6% reduction. Grain protein concentration increased under severe HS in all three genotypes, with the largest increase in the diploid genotype (24%). This increase was probably influenced by a concentration effect caused by reduced starch deposition, although increased synthesis of stress-response proteins has also been reported in cereals (Dupont et al. 2006).

The response profiles may partly arise from variation in genome composition. Wheat subgenomes are functionally asymmetric in their contributions to developmental and stress-responsive traits (Feldman et al. 2012). The lineages represented here also differ in their domestication trajectories, breeding histories, and levels of genetic diversity (Haudry et al. 2007; Borrill et al. 2019). Thus, the observed contrasts may reflect genome constitution as well as lineage-specific selection.

### 4.2 Transcriptomic Responses to HS in Diploid, Tetraploid, and Hexaploid Wheat Genotypes

We conducted a transcriptomic analysis to investigate HS responses in the three wheat genotypes. The diploid, tetraploid, and hexaploid analyses identified 859, 757, and 2,227 DEGs. Raw expressed-gene counts are not directly comparable across ploidy levels: the 39,847 expressed genes in the tetraploid genotype corresponded to 22,092 nonredundant gene units after homoeolog grouping, whereas the 68,303 genes in the hexaploid genotype corresponded to 26,268 units. Within the heat-responsive sets, the tetraploid and hexaploid DEGs represented 562 and 1,445 homoeolog groups, respectively. These values fall within the range reported in previous wheat HS studies (Arenas-M et al. 2021; Paul et al. 2022; Kumar et al. 2015; Duan et al. 2025; Kalaipandian et al. 2023).

Despite quantitative differences in the DEG sets, a common HS response was observed across all three genotypes. This response included upregulation of genes encoding heat shock proteins (HSPs), particularly members of the HSP70 and HSP90 families (Fig. 3B), which participate in protection against protein denaturation and aggregation (Takahagi et al. 2018; Lu et al. 2020). GO terms related to “response to heat” and “protein folding” (group I, Fig. 4A) were significantly enriched among upregulated genes in all three analyses. These results support a shared chaperone-centered response in the examined wheat genotypes (Lu et al. 2020; Zheng et al. 2025).

In addition to this common response, the hexaploid genotype had several enriched pathways that were not detected in the other two analyses (group IV, Fig. 4A). Enrichment of “protein quality control via the ERAD pathway” indicates overrepresentation of genes assigned to ERAD and protein-quality-control functions among the upregulated genes. ERAD removes misfolded proteins from the endoplasmic reticulum through recognition, ubiquitination, dislocation, and degradation, contributing to ER homeostasis (Reyes-Impellizzeri and Moreno 2021). Because GO enrichment does not directly measure pathway activity, these data support heat-responsive regulation of protein-quality-control genes rather than demonstrating enhanced ERAD function.

Enrichment of “mRNA processing” and “cis splicing via spliceosome” in hexaploid wheat indicates that genes annotated for RNA metabolism were overrepresented among its upregulated genes. AS can increase proteome diversity and regulate transcript abundance under stress (Alhabsi et al. 2025). In wheat, alternative splicing of heat shock transcription factors can generate transcripts that differ in their capacity to induce heat shock protein genes and affect thermotolerance (Wen et al. 2023). Combined with the event-level analysis below, the enrichment results support heat-associated remodeling of RNA processing in the examined hexaploid genotype.

Among downregulated genes in hexaploid wheat, “starch biosynthetic process” was enriched (group III, Fig. 4B). This expression pattern is consistent with the lower final starch concentration under HS and with previous results in durum wheat (Arenas-M et al. 2021). Because transcriptome sampling preceded final grain analysis, the temporal ordering is compatible with a relationship between early transcriptional repression and reduced starch accumulation, but it does not establish causality.

A total of 757 DEGs were detected in the tetraploid genotype, with two nonredundant GO terms enriched among the upregulated genes (Fig. 4A). Its proteostasis-related DEGs included *Fes1C*, which encodes an HSP70 nucleotide-exchange factor, and *CLPB3*, which encodes a caseinolytic peptidase involved in protein disaggregation (Lu et al. 2022). Their differential expression documents heat-responsive regulation of genes associated with HSP70 cycling and protein disaggregation in the tetraploid genotype, without by itself establishing changes in the corresponding protein activities. The diploid genotype, which had a 59.4% mean yield reduction under TT2 in Exp. 2, had 859 DEGs. Its upregulated gene set shared several GO terms with the hexaploid genotype (group II, Fig. 4A), including responses to salt, hypoxia, and heavy metals. These enrichments indicate that several abiotic-stress response categories were represented in the diploid DEG set.

Together, the transcriptomic results define genotype-associated differences in both the scale and functional composition of the HS response. Because raw expressed-gene counts increase with genome complexity, homoeolog grouping provides the more informative comparison across the three analyses; after grouping, the heat-responsive sets still differed substantially in size and composition. These patterns are consistent with reported molecular differences among modern hexaploid wheat and its diploid and tetraploid relatives (Li et al. 2018).

### 4.3 HS Elicits Distinct Alternative Splicing Responses in Wheat

Beyond transcription, the results identify AS as a component of the wheat HS response. AS increases proteome diversity and contributes to plant responses to environmental stress, including heat (Alhabsi et al. 2025). The tetraploid genotype had the largest number of significant AS events. GO enrichment among genes containing differential splicing events yielded one term in the diploid and six terms in the hexaploid genotype, whereas no term was detected in the tetraploid genotype. Event counts and functional enrichment therefore describe different properties of the AS response.

The relationship between polyploidy and AS is complex. Functional redundancy among homoeologs might reduce selective pressure on splice variants (Yu et al. 2020), whereas other studies have linked polyploidy to stress resilience mediated in part by AS (Li et al. 2024). Wheat subgenomes can also differ in AS during development and stress (Yu et al. 2020; Li et al. 2024). In the present study, the three A-, B-, and D-subgenome *NF-YB* homoeologs in the hexaploid genotype showed concordant heat-induced splicing changes, whereas corresponding changes were not detected in the diploid and tetraploid orthologs. This locus-level concordance identifies *NF-YB* as a candidate for functional analysis.

To examine this AS event in detail, we analyzed the *Nuclear Factor Y subunit B* (*NF-YB* ) gene. This gene showed significant heat-induced AS (|ΔPSI| *>* 0.1, FDR *<* 0.05) across the A-, B-, and D-subgenome homoeologs of the hexaploid genotype; a corresponding change was not detected in the diploid and tetraploid orthologs. NF-YB is a component of the conserved heterotrimeric NF-Y transcription factor complex (NF-YA/NF-YB/NF-YC), which binds CCAAT-box motifs in target promoters and regulates gene expression (Zhao et al. 2016). The NF-YB histone-fold domain (HFD) is required for heterodimerization with NF-YC, stabilization of the complex, and DNA binding by NF-YA (Zhao et al. 2016). NF-Y complexes participate in plant development and responses to drought, salinity, and temperature (Zhao et al. 2016). In rice, temperature-dependent interactions within an NF-Y complex regulate *QT12* and affect grain thermotolerance (Li et al. 2025); in Arabidopsis, AtNF-YB3 contributes to heat tolerance through heat-responsive gene expression (Sato et al. 2019). These studies provide a biological basis for functional analysis of the wheat splice variant.

The E6-containing transcript model *TraesCS1D03G0689500.1* is predicted to encode a full-length NF-YB protein. Because E6 spans 51 bp, its removal is in frame, and the E6-skipped model *TraesCS1D03G0689500.2* is predicted to encode a protein shorter by the 17-residue sequence MGQQVAYNPGMVYMQPQ. This segment lies adjacent to the conserved HFD, a position from which effects on NF-YB/NF-YC assembly, DNA binding or target selection are mechanistically plausible. The E6-skipped transcript may therefore encode a dominant-negative regulator or a functional variant with altered specificity, providing a concrete model for protein-expression, complex-assembly, DNA-binding, and thermotolerance assays.

## 5 Conclusions

Short-term HS during early grain filling elicited distinct phenotypic and molecular responses in the examined diploid, tetraploid, and hexaploid wheat genotypes. Integrating yield traits, differential expression, GO enrichment, and alternative splicing showed that the heat responses differed not only in magnitude but also in the biological processes represented among responsive genes and splice events. In the hexaploid genotype, concordant heat-induced E6 skipping across the three *NF-YB* homoeologs, supported by RT-PCR and sequence confirmation for the D-genome locus, defines *NF-YB* splicing as a specific, experimentally supported candidate response. These results provide a multilevel framework for prioritizing regulatory candidates associated with wheat grain responses to heat.

**Electronic supplementary material. Online Resource 1 (PDF): Supplementary Fig. S1. Meteorological conditions during the wheat growing seasons.** Daily minimum and maximum temperature, precipitation, and incident photosynthetically active radiation at the experimental site during Experiments 1 and 2, with the timing of developmental stages and management activities.

**Online Resource 2 (PDF): Supplementary Fig. S2. RT-qPCR validation of RNA-seq expression patterns.** Relative expression of *HSP17.6II* and *HSP26* under CT, TT1, and TT2 in the three examined genotypes.

Online Resource 3 (PDF): Supplementary Fig. S3. Sanger-sequence confirmation of *NF-YB* splice variants. Chromatograms and sequence alignment of the E6-included and E6-skipped transcripts in *T. aestivum*.

Online Resource 4 (PDF): Supplementary Fig. S4. Quantification of heat-induced exon E6 skipping in *NF-YB*. E6 inclusion calculated from RT-PCR band intensities in three paired biological replicates.

**Online Resource 5 (PDF): Supplementary Table S1. Measured temperatures and heat loads during the thermal treatments.** Four-day mean temperatures at spike height and cumulative degree days above 18*^◦^*C for each genotype and treatment.

**Online Resource 6 (PDF): Supplementary Table S2. Thermal-treatment effects on grain dimensions and growth dynamics.** Individual grain weight, grain dimensions, and grain-filling rate and duration in Experiments 1 and 2.

**Online Resource 7 (XLSX): Supplementary Table S3. Differentially expressed genes under HS.** Gene identifiers, fold changes, adjusted *P* -values, and annotations for the three genome-specific analyses.

**Online Resource 8 (XLSX): Supplementary Table S4. GO enrichment of heat-responsive genes.** Biological-process terms enriched among upregulated and downregulated genes, with Benjamini–Hochberg adjustment and rrvgo semantic reduction.

**Online Resource 9 (XLSX): Supplementary Table S5. Differential alternative-splicing events under HS.** Whippet/betAS event-level results for the three examined genotypes.

Online Resource 10 (XLSX): Supplementary Table S6. *NF-YB* alternative splicing across *T. aestivum* homoeologs. PSI values, ΔPSI, and FDR for the A-, B-, and D-subgenome copies.

**Online Resource 11 (XLSX): Supplementary Table S7. Oligonucleotide primers.** Primers used for RT-qPCR, RT-PCR, cloning, and sequence confirmation. **Online Resource 12 (XLSX): Supplementary Table S8. RNA-seq quality and pseudoalignment statistics.** Read-quality metrics and Kallisto pseudoalignment rates for the 18 libraries.

## Author contributions

AAM conducted experiments, performed formal analysis, and drafted the manuscript. IM and DL conducted experiments and curated data. CU contributed to study conception and revised the manuscript. DFC conceived the study, supervised the research, acquired funding, and revised the manuscript. JC conceived and supervised the study, administered the project, and revised the manuscript. All authors read and approved the final manuscript.

## Declarations

### Data availability

Raw RNA-seq reads are available in NCBI Gene Expression Omnibus under accession GSE315971. Additional data supporting the reported results are provided in the online resources.

