## Supplementary material for "Multilevel Characterization Reveals Divergent Heat Responses Among Diploid, Tetraploid, and Hexaploid Wheat Genotypes": ESM_1_bioRxiv.pdf

### Electronic Supplementary Material 1

**Journal:** *bioRxiv*

---

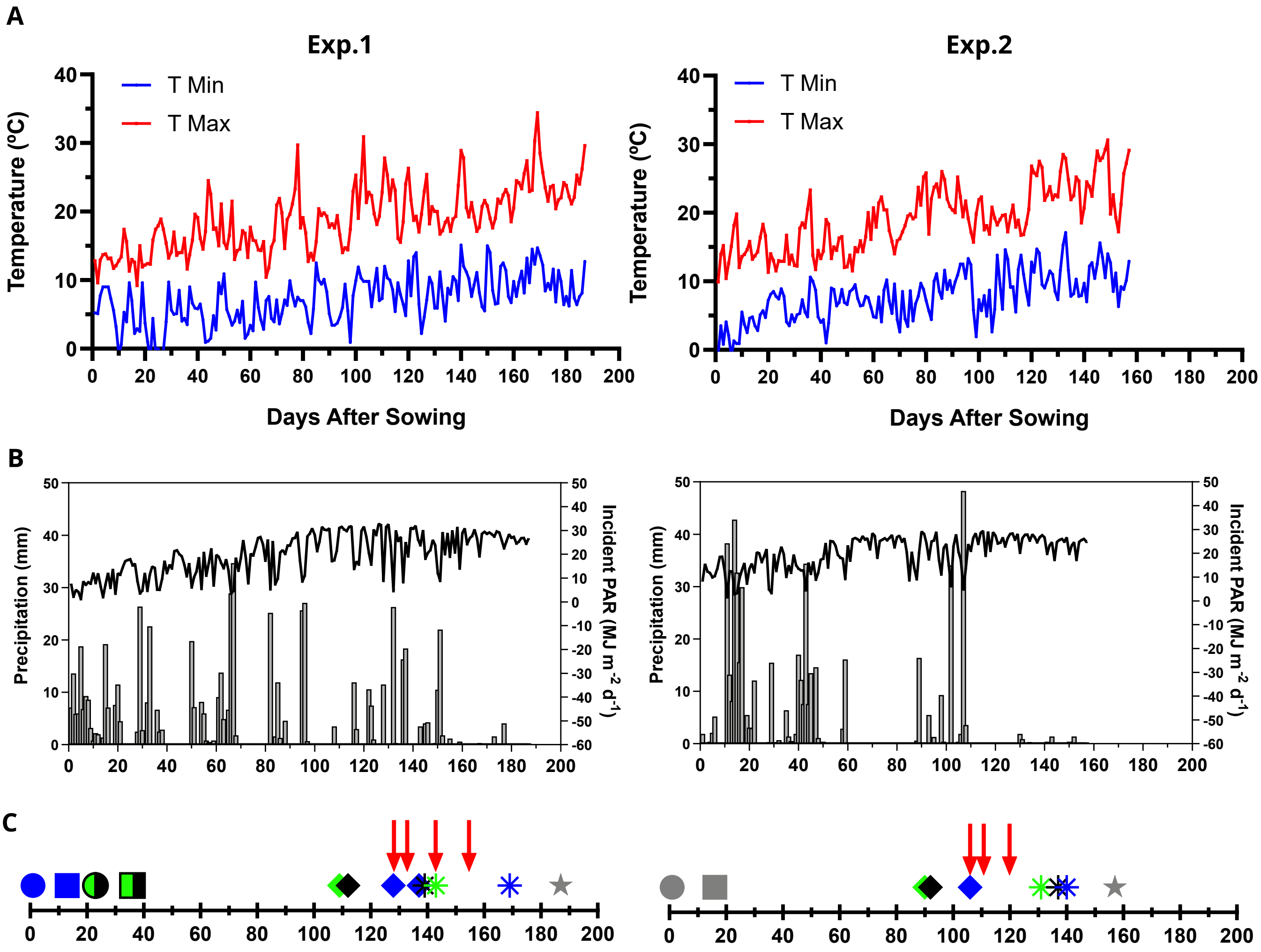


**Supplementary Fig. S1 Meteorological conditions during the wheat growing seasons.** Daily minimum and maximum temperature, precipitation, and incident photosynthetically active radiation at the experimental site during Experiments 1 and 2, with the timing of developmental stages and management activities.
