## Supplementary material for "Multilevel Characterization Reveals Divergent Heat Responses Among Diploid, Tetraploid, and Hexaploid Wheat Genotypes": ESM_2_bioRxiv.pdf

### Electronic Supplementary Material 2

**Journal:** *bioRxiv*

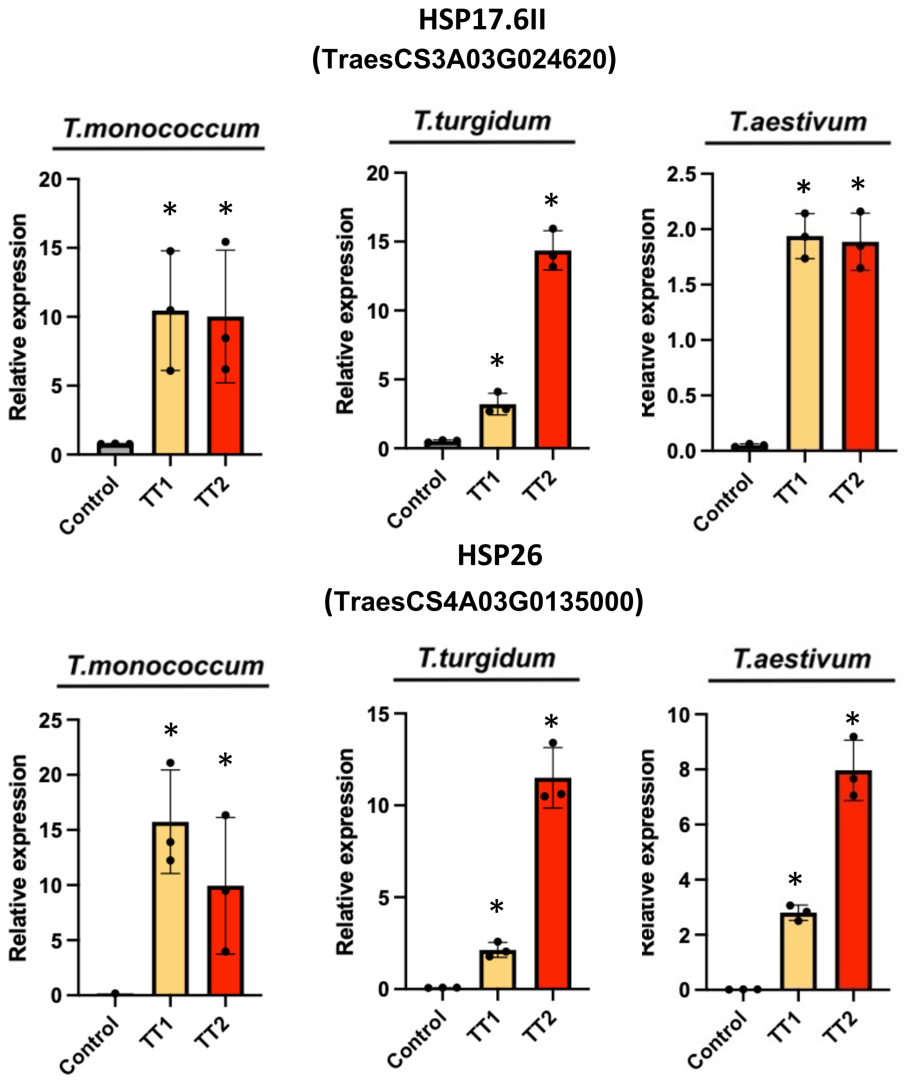


**Supplementary Fig. S2 RT-qPCR validation of RNA-seq expression patterns.** Relative expression of *HSP17.6II* and *HSP26* under CT, TT1, and TT2 in the three examined genotypes.
