## Supplementary material for "Multilevel Characterization Reveals Divergent Heat Responses Among Diploid, Tetraploid, and Hexaploid Wheat Genotypes": ESM_3_bioRxiv.pdf

### Electronic Supplementary Material 3

**Journal:** *bioRxiv*

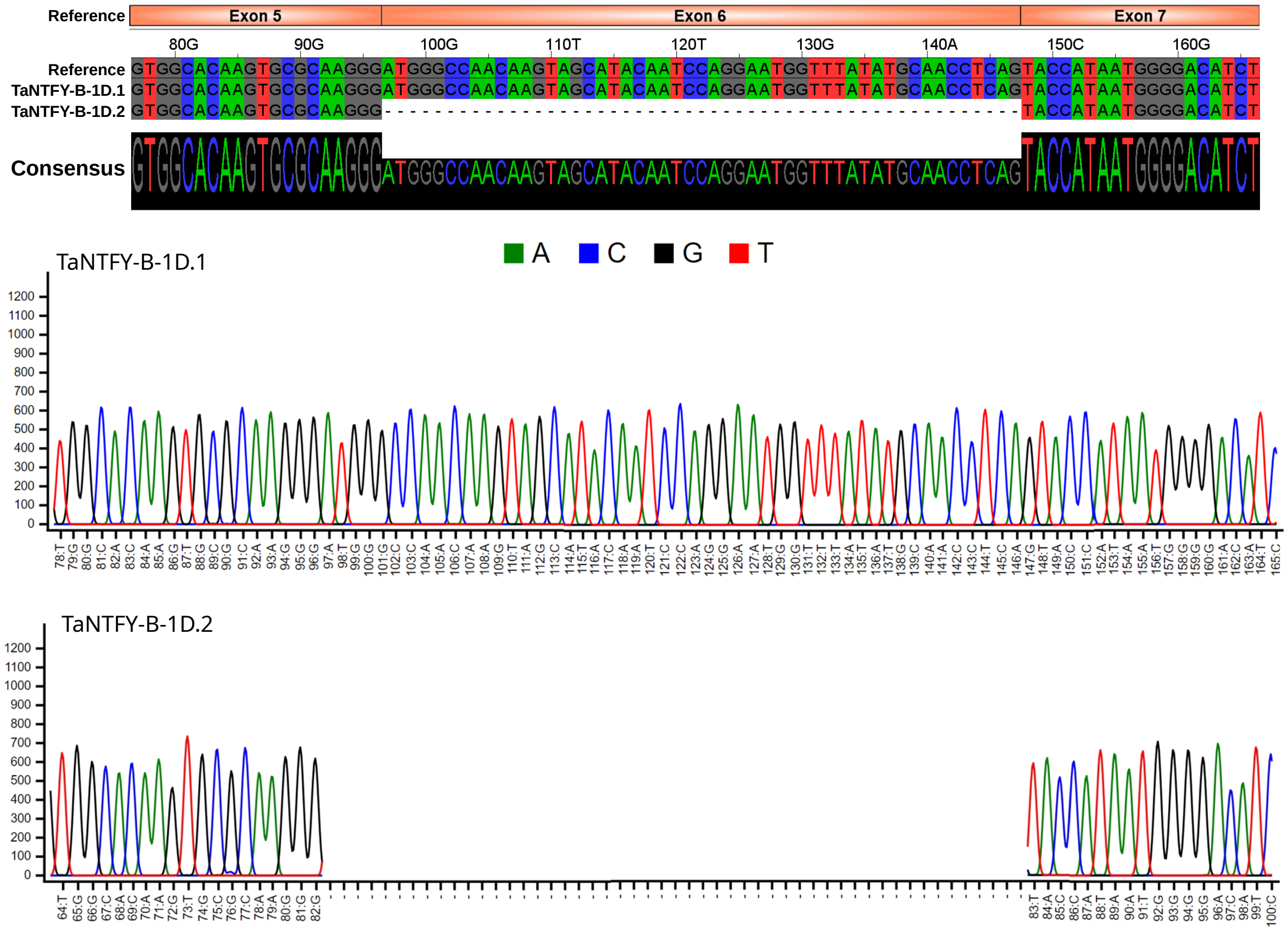


**Supplementary Fig. S3 Sanger-sequence confirmation of *NF-YB* splice variants.** Chromatograms and sequence alignment of the E6-included and E6-skipped transcripts in *T. aestivum*.
