## Supplementary material for "Multilevel Characterization Reveals Divergent Heat Responses Among Diploid, Tetraploid, and Hexaploid Wheat Genotypes": ESM_4_bioRxiv.pdf

### Electronic Supplementary Material 4

**Journal:** *bioRxiv*

---

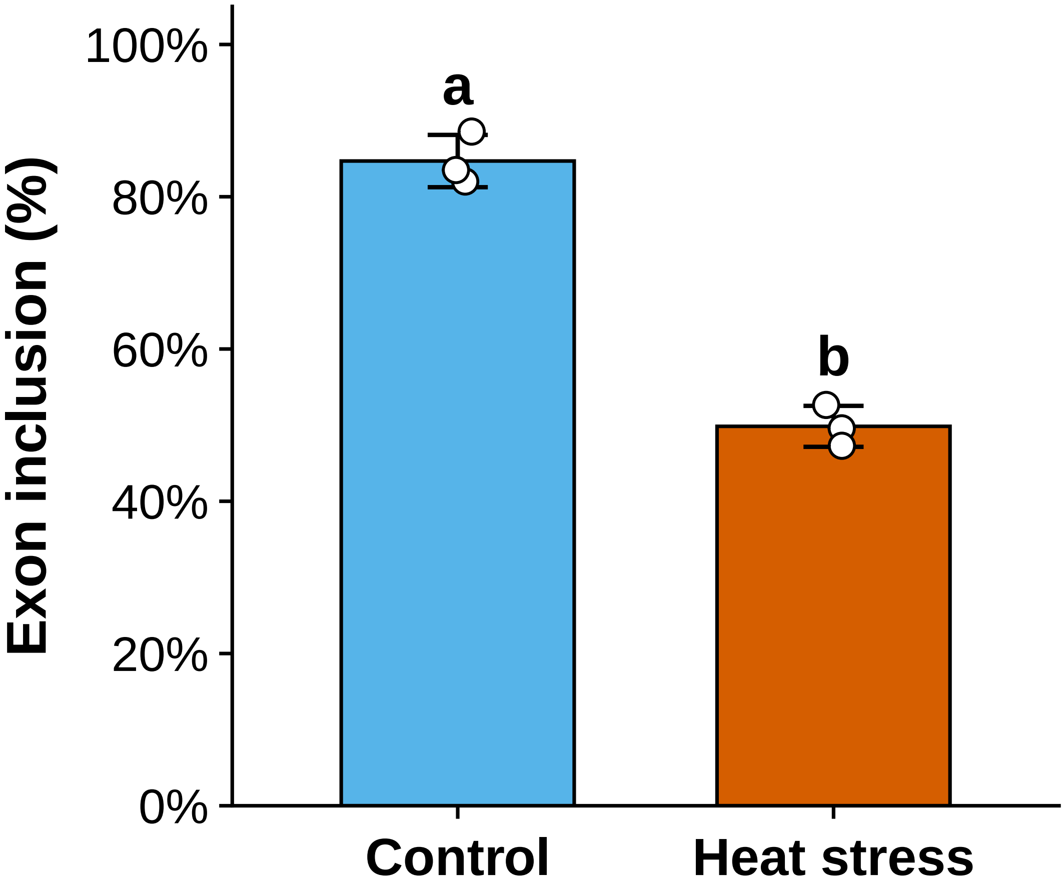


**Supplementary Fig. S4 Quantification of heat-induced exon E6 skipping in *NF-YB*.** E6 inclusion calculated from RT-PCR band intensities in three paired biological replicates.
