## Supplementary material for "Multilevel Characterization Reveals Divergent Heat Responses Among Diploid, Tetraploid, and Hexaploid Wheat Genotypes": ESM_5_bioRxiv.pdf

### Electronic Supplementary Material 5

**Journal:** *bioRxiv*

**Supplementary Table S1 Measured temperatures and heat loads during the thermal treatments.** Four-day mean temperatures at spike height and cumulative degree days above 18°C for each genotype and treatment.

| | | | Mean temperature (°C) | | | Heat load (°C days > 18°C) | |
|---|---|---|---|---|---|---|---|
| Experiment | Ploidy | Genotype | CT | TT1 | TT2 | TT1 | TT2 |
| 1 | Diploid | Mon1 | 19.5 | 27.4 | – | 37.6 | – |
| 1 | Diploid | Mon8 | 19.5 | 27.2 | – | 36.8 | – |
| 1 | Tetraploid | Corcolen | 17.0 | 24.0 | – | 24.0 | – |
| 1 | Tetraploid | Queule | 17.0 | 23.9 | – | 23.6 | – |
| 1 | Hexaploid | Impulso | 13.8 | 22.7 | – | 18.8 | – |
| 1 | Hexaploid | Fritz | 13.8 | 22.8 | – | 19.2 | – |
| 2 | Diploid | Mon8 | 14.9 | 26.3 | 33.7 | 33.2 | 62.8 |
| 2 | Tetraploid | Queule | 20.2 | 28.5 | 33.7 | 42.0 | 62.8 |
| 2 | Hexaploid | Fritz | 12.6 | 27.6 | 31.2 | 38.4 | 52.8 |

CT, ambient control; TT1, moderate heating; TT2, severe heating. Heat load is cumulative degree days above 18°C during the four-day treatment. A dash indicates that TT2 was not included in Experiment 1.
