## Supplementary material for "Multilevel Characterization Reveals Divergent Heat Responses Among Diploid, Tetraploid, and Hexaploid Wheat Genotypes": ESM_6_bioRxiv.pdf

**Electronic Supplementary Material** **Journal:** *bioRxiv*

**Supplementary Table S2 Thermal-treatment effects on grain dimensions and growth dynamics.** Individual grain weight, grain dimensions, and grain-filling rate and duration in Experiments 1 and 2.

| | | | | | Grain dimensions | | | Grain growth dynamics | | |
|---|---|---|---|---|---|---|---|---|---|---|
| Exp. | Ploidy | Genotype | Treatment | IGW-G1 (mg) | Length (mm) | Width (mm) | Area ($mm^2$) | Calculated IGW-G1 (mg) | Filling rate (mg $day^{-1}$) | Filling duration (days) |
| 1 | Diploid | Mon1 | CT | 35.3 | 7.4 | 3.1 | 15.8 | 32.6 | 0.9 | 42.5 |
| | | | TT1 | 29.0 | 7.2 | 2.9 | 14.3 | 30.0 | 0.9 | 38.7 |
| | | Mon8 | CT | 35.3 | 8.2 | 3.3 | 18.1 | 33.1 | 0.9 | 42.9 |
| | | | TT1 | 29.8 | 8.2 | 3.2 | 17.4 | 34.8 | 0.9 | 43.0 |
| | Tetraploid | Corcolen | CT | 84.4 | 8.1 | 3.9 | 23.4 | 72.4 | 1.6 | 47.3 |
| | | | TT1 | 82.0 | 8.2 | 3.9 | 23.5 | 71.7 | 1.6 | 47.1 |
| | | Queule | CT | 78.8 | 8.1 | 3.8 | 22.5 | 67.6 | 1.6 | 44.8 |
| | | | TT1 | 76.5 | 7.9 | 3.6 | 21.3 | 63.5 | 1.6 | 42.2 |
| | Hexaploid | Impulso | CT | 59.5 | 6.7 | 3.9 | 19.5 | 55.0 | 1.3 | 44.3 |
| | | | TT1 | 61.6 | 6.9 | 3.9 | 19.9 | 52.9 | 1.2 | 44.9 |
| | | Fritz | CT | 89.0 | 7.9 | 4.3 | 26.0 | 86.4 | 1.7 | 53.5 |
| | | | TT1 | 84.3 | 8.0 | 4.3 | 26.4 | 76.9 | 1.8 | 45.7 |
| | SEM | | | 6.8 | 0.2 | 0.1 | 1.1 | 5.7 | 0.1 | 1.0 |
| | Genotype | | | **** | **** | **** | **** | **** | **** | *** |
| | Treatment | | | ns | ns | * | ns | ** | ns | ns |
| | Genotype × Treatment | | | ns | ns | ns | ns | ns | ns | ns |
| 2 | Diploid | Mon8 | CT | 35.2 | 7.3 | 3.1 | 15.2 | 30.7 | 1.0 | 36.8 |
| | | | TT1 | 23.7 | 6.8 | 2.7 | 12.6 | 25.1 | 1.0 | 31.8 |
| | | | TT2 | 18.7 | 6.6 | 2.6 | 12.1 | 21.8 | 1.2 | 28.0 |
| | Tetraploid | Queule | CT | 79.7 | 8.2 | 3.8 | 23.3 | 76.2 | 1.7 | 46.7 |
| | | | TT1 | 73.5 | 8.2 | 3.6 | 22.1 | 65.5 | 1.3 | 48.0 |
| | | | TT2 | 65.1 | 8.0 | 3.6 | 21.1 | 61.0 | 1.3 | 46.2 |
| | Hexaploid | Fritz | CT | 79.6 | 7.8 | 4.4 | 25.2 | 77.7 | 1.8 | 49.1 |
| | | | TT1 | 70.4 | 7.7 | 4.2 | 23.9 | 62.1 | 1.6 | 44.6 |
| | | | TT2 | 62.7 | 7.8 | 4.2 | 24.2 | 58.1 | 1.5 | 43.0 |
| | SEM | | | 7.9 | 0.2 | 0.2 | 1.7 | 7.2 | 0.1 | 2.5 |
| | Genotype | | | **** | **** | **** | **** | **** | **** | **** |
| | Treatment | | | **** | ** | ** | ** | **** | ns | ** |
| | Genotype × Treatment | | | ns | * | ns | ns | ns | ns | ns |

IGW-G1, individual grain weight at position G1. Calculated IGW-G1 is the fitted individual grain weight at the end of grain growth. Values are means of three plot replicates. SEM, standard error of the mean; ns, not significant; * $P < 0.05$; ** $P < 0.01$; *** $P < 0.001$; **** $P < 0.0001$.
